# Automatic Single-Cell Discrimination by Cellular Appearance using Convolutional Neural Network

**DOI:** 10.1101/446823

**Authors:** Yoshikazu Matsuoka, Ryusuke Nakatsuka, Tatsuya Fujioka

## Abstract

Morphological images of cells contain extensive information, which help biologists to infer the type and state of cells to some degree based on their morphology. Convolutional Neural Network, a neural network architecture, is a powerful tool used for image recognition. However, whether it can be used to classify cells on the basis of their morphology remains unclear. In this study, we demonstrate that 10 different hematopoietic tumor cell lines with similar morphologies that biologists find difficult to distinguish can be classified with >90% accuracy using only their bright-field images when analyzed using a Convolutional Neural Network. This novel and simple system using bright-field images of cells could be a powerful analytical tool for cell type discrimination and could also be applied to the clinical diagnoses of hematopoietic tumors.

## Introduction

Cell categorization is essential for elucidating the function of each cell population. This task is usually undertaken by detecting the expression of marker genes/proteins under a fluorescence microscope or by flow cytometry. These florescent based analysis tools usually can use up to 12 colors by their limitation of fluorescence overlap (Autissier et al., 2010), though improvements for increasing the number of usable channels are still underway. Cytometry using time-of-flight (CyTOF) considers large number (up to 42) of parameters compared with conventional flow cytometry analysis, and thus it can precisely analyze the details of cell populations using intracellular and extracellular markers (Bendall et al., 2011; Kay et al., 2016). In addition, gene expression data obtained by single-cell RNA-seq is also useful for cell-classification (Han et al., 2018; Treutlein et al., 2014).

Alternatively, morphological observation of cells using a microscope is widely performed in the field of cell biology. We can obtain extensive information regarding cell features from their morphology; thus, biologists can then infer the type and state of cells with a certain degree of accuracy by comparing the images obtained from microscopic observations and standard morphological findings for cells. However, it is difficult to discriminate the cells with similar morphologies using only cell bright-field image.

Currently, machine learning (ML) is used for data classification, for instance, in gene expression data profiling (Singh et al., 2016; Alipanahi et al., 2015). Moreover, Convolutional Neural Network (CNN), a neural network architecture, is an extremely potent tool in the field of image classification (Krizhevsky et al., 2012; Simonyan and Zisserman, 2014), and it has already been employed in many generally usable web applications, including Google, Facebook, and Twitter (LeCun et al., 2015).

The concept of image-based cell classification using ML has already been conceived by many researchers and is expected to be applied to cell biological analysis (Grys et al., 2017; Kan, 2017; Caicedo et al., 2017; Doan M et al., 2018). However, there is almost no report presenting experimental data of discrimination accuracy of cells with similar morphology using bright-field image based ML. To build a high-quality model for discrimination on CNN, it is required to obtain a large quantity of high quality images (Bengio et al., 2013). In this respect, imaging flow cytometer is very useful for obtaining a large quantity of images for CNN (Han et al., 2016). Additionally, using imaging flow cytometer, we can easily obtain over thousand dimensional data even from grayscale digital image of single-cell.

In this study, we combined imaging flow cytometry (IFC) analysis and CNN to classify different populations of cells using their morphological features. As a result, the classification accuracy of human hematopoietic tumor-derived cell lines with similar morphology using image-based ML was greatly exceeded compared with classification accuracy by biologists. This indicates that the image-based cell discrimination system is useful for cell biological research.

## Results and discussion

### Human hematopoietic tumor-derived cell lines could be discriminated based on their bright-field images using CNN

First, we collected bright-field images of 10 different human hematopoietic tumor cell lines (including acute myeloid leukemia, chronic myeloid leukemia, B-cell acute lymphoblastic leukemia, and myeloma; listed in Table S1) using IFC (90,000 images per group) (Figures 1 and S1). Each pixel of these digital images of cells was defined by a numerical value from 0 to 255 (8-bit, grayscale) reflecting its brightness (48 × 48 pixels/image) (Figure 2A). Therefore, we tested whether these multi-dimensional data could be used for cell-classification using CNN. We prepared training and test data (Figure 3A), and then, extracted features from training images obtained for each cell line and train a model (deposited in https://figshare.com/s/0ff584cb070cd03164aa) for automatic discrimination by CNN (Figure 3B). The accuracy of classification for each cell line was increased according to the increasing number of images for training (Figures 2B and C). As shown in Figure 2B, the classification accuracy for each cell line gradually increased until epoch (the number of iterations of passes through the entire training set) 40. Finally, the results from this system achieved over 90% accuracy for discrimination of the 10 hematopoietic tumor cell lines (Figures 2C and D). These results suggest that each bright-field image contains a sufficient amount of information to distinguish each cell type.

**Figure 1.**
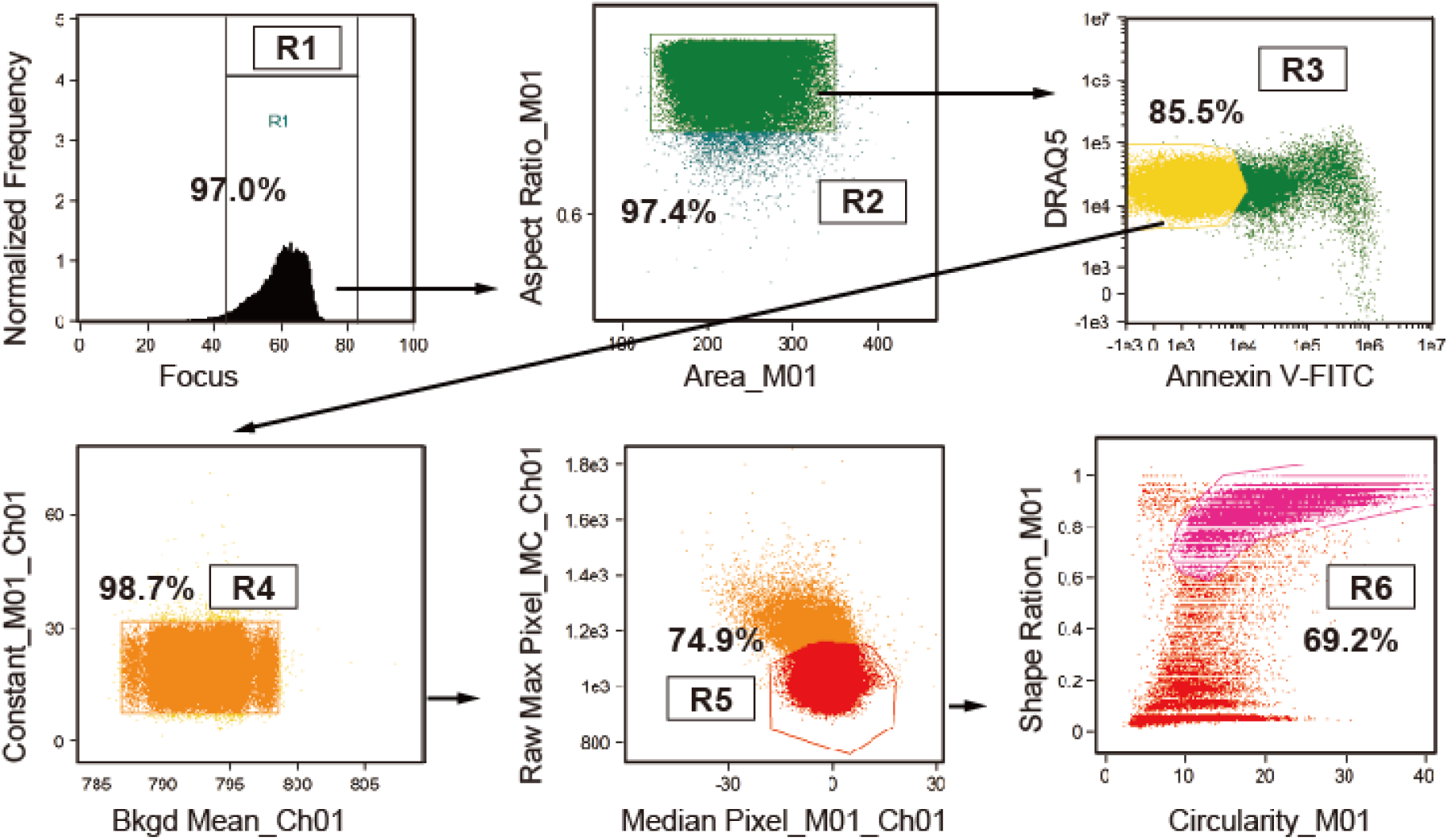
**Gating strategies for IFC for obtaining the cell images**. An R1 gate was set on the focused cells. The doublet cells in the low aspect ratio area were excluded by the R2 gate. Next, DRAQ5-positive and Annexin V-negative nucleated cells were gated (R3). Then, the cells located near the center of the image field and round cells were selected by R4 to R6 gating.

**Figure 2.**
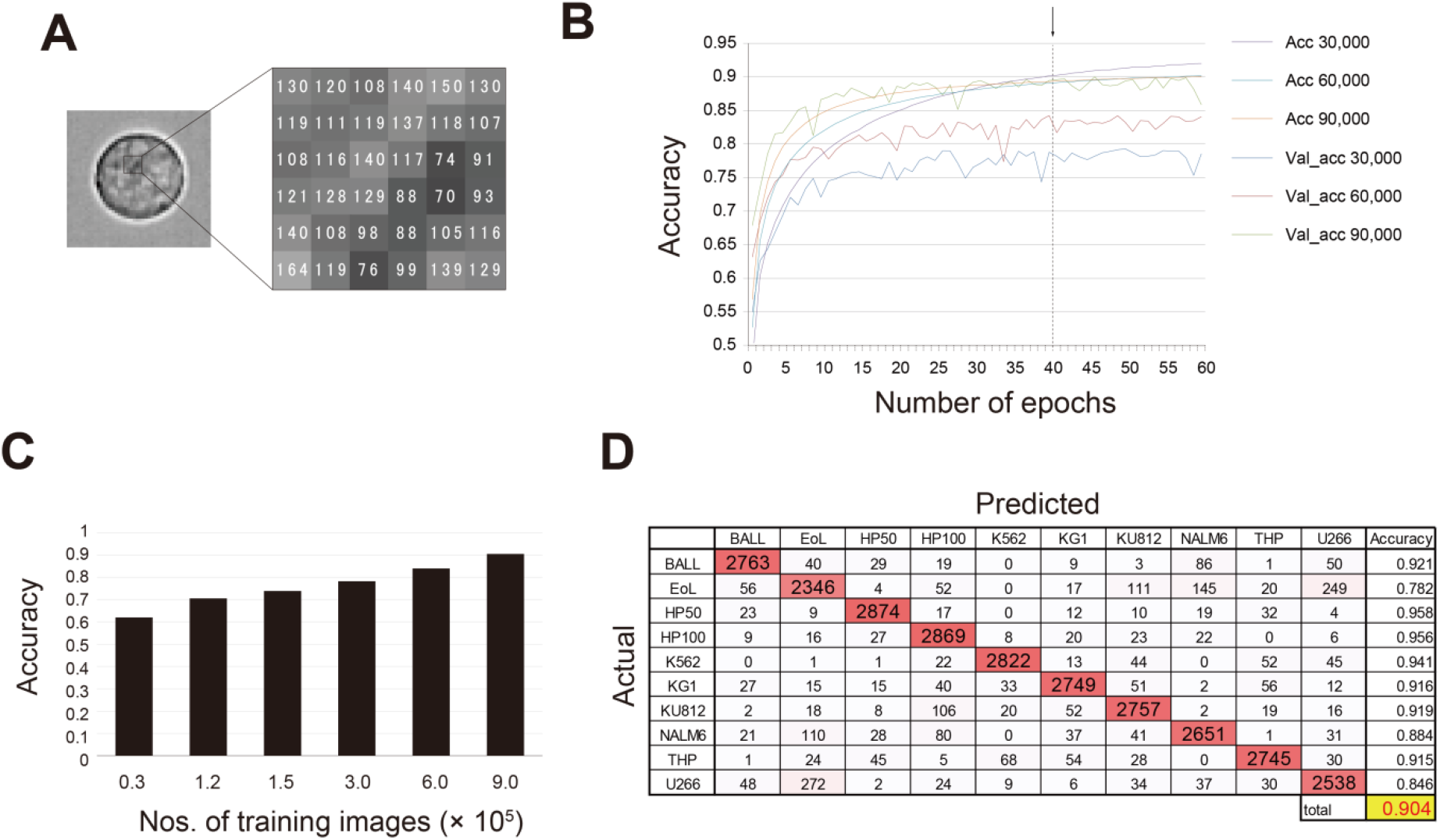
**Classification of 10 kinds of hematopoietic tumor cell lines by CNN** (A) A schematic image of numerical components of a digital image. The cell image was obtained by IFC using 40× objective lens. The representative image of K562 cell is provided. The numerical data in 6 × 6 pixel regions are magnified. The value (0 to 255, 8-bit grayscale image) of digital data in each pixel reflects its brightness. (B) Fine-tuning of the epoch numbers of CNN. Arrow and dotted line indicate the peak of classification accuracy (epoch 40) for 10 discrimination classes. Acc: Accuracy of training data. Val_acc: Accuracy of test data. The numbers after Acc or Val_acc represent the numbers of training data/group. The data indicate the mean value obtained from three runs of CNN using the same training and test data. (C) A comparison of the accuracy among the numbers of training data. The data were obtained at epoch 40. The data indicate the mean value obtained from three runs of CNN using the same training and test data. (D) Confusion matrix of the results of classification by CNN. The data in the table are colored a darker red with increasing value. The data are representative from three runs of CNN.

**Figure 3.**
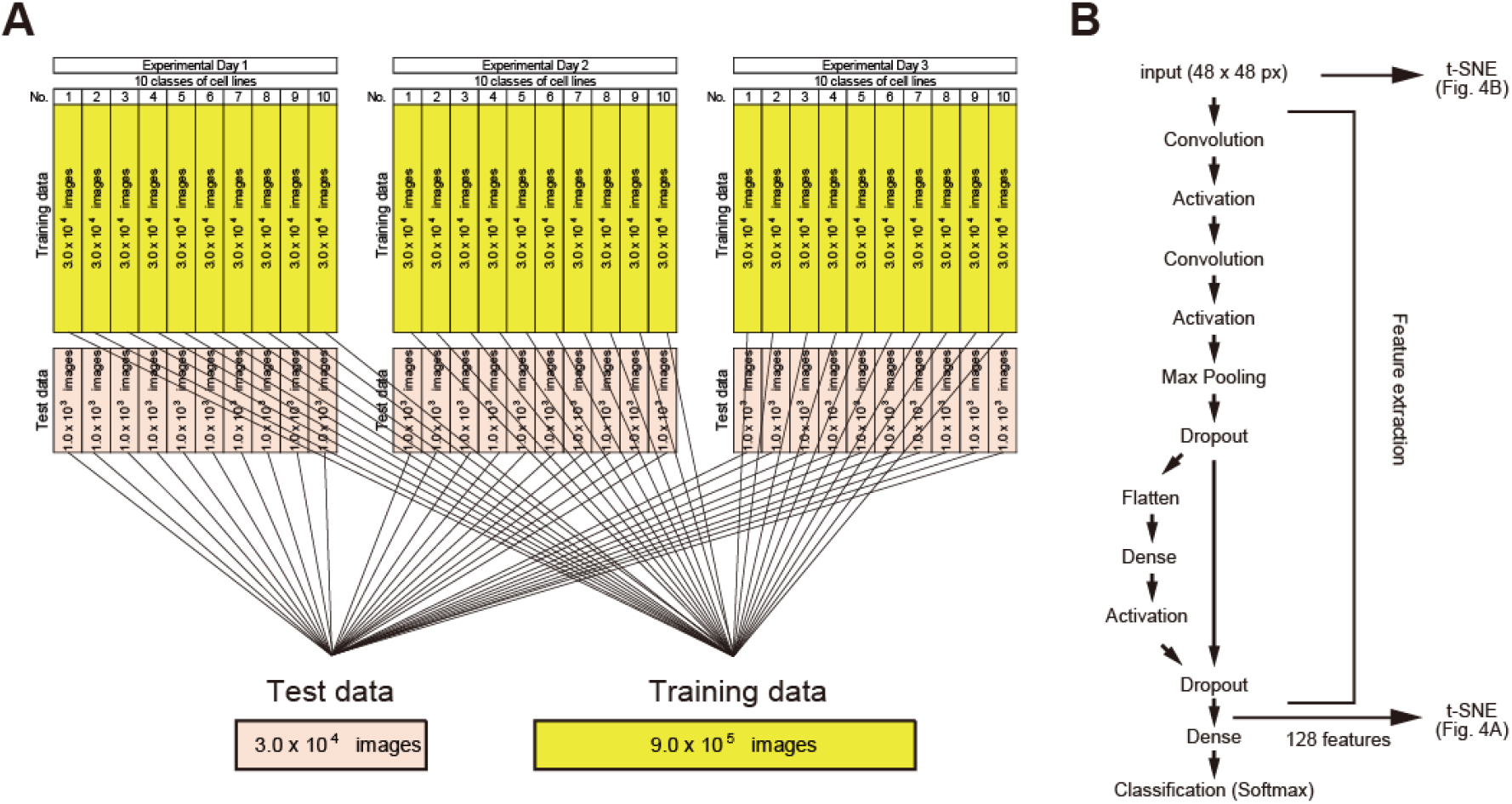
**Schematic illustration of preparation of training and test data for CNN and the construction of CNN used in this study** (A) The data obtained from three independent experimental days were combined and used for CNN. (B) The signal of the cell bright-field images (48 × 48 pixels/image, 9.0 × 10^4^ images/group, a total of 9.0 × 10^5^ images) captured by IFC were input into CNN. After feature extraction, the features of 3,000 cells/group were exported and compressed into 2D data by t-SNE (Figure 4A). The raw numerical data of the same cells (3,000 cells/group) were also compressed into 2D by t-SNE (Figure 4B).

Among the 10 hematopoietic tumor cell lines, and regardless of hematopoietic tumor type, it seemed that some could be accurately classified using the CNN algorithm while some could not (Figure 2D). Therefore, we attempted to separately discriminate three different cell lines with good (HP50-2, HP100-1, and K562) or poor classification accuracy (EoL-1, NALM-6, and U266). The results demonstrated that the classification accuracy for the former group was about 97%, whereas that for the latter group was about 84% (Table 1). This revealed that some of the cell lines were easy and others were difficult to distinguish using CNN.

**Table 1.** **The discrimination accuracy for cell lines with good and poor classification rates**.

|  |  | Predicted |  |  |  |
| --- | --- | --- | --- | --- | --- |
| Actual | Good_3 | HP50 | HP100 | K562 | Accuracy |
|  | HP50 | 2888 | 112 | 0 | 0.963 |
|  | HP100 | 132 | 2846 | 22 | 0.949 |
|  | K562 | 2 | 35 | 2963 | 0.988 |
|  |  |  |  | Total | 0.966 |
| Actual | Poor_3 | EoL | NALM | U266 | Accuracy |
|  | EoL | 2275 | 333 | 392 | 0.758 |
|  | NALM | 169 | 2785 | 46 | 0.928 |
|  | U266 | 451 | 93 | 2456 | 0.819 |
|  |  |  |  | Total | 0.835 |

### The classification accuracy of cells by CNN were far superior to those by biologists

Next, we examined whether the classification accuracy of these cell lines using this system was superior or inferior to that performed by biologists. The results revealed that biologists found it difficult to identify these cell lines using only bright-field images (success rates 8%–35%) (Table S2). In addition, it took approximately three hours to train a model for CNN using 9.0 × 10^5^ cells; however, it took ≤10 s to discriminate 3.0 × 10^4^ cells using the trained data. Alternatively, it took approximately 30 minutes to 1 hour to discriminate 100 cells by biologists. Even after repeated training, the accuracy with which biologists could classify these cell lines did not improve at all (Table S2).

To elucidate why this system can discriminate cell types with such high accuracy, the features extracted from each single-cell were compressed into two dimension (2D) by t-Distributed Stochastic Neighbor Embedding (t-SNE) (van der Maaten and Hinton 2018). In the 2D plot produced by this approach, the extracted features of 10 different hematopoietic tumor cell lines were segregated (Figure 4A). Furthermore, the raw pixel data of images of these cell lines were mixed with each other, and they were inseparable (Figure 4B). The extracted features of EoL and U266, which were difficult to discriminate from each other by CNN, were partially overlapped. Hence, the accuracy of classification between these cell lines was lower compared with that for the other cell lines, as shown in Figure 2D and Table 1.

**Figure 4.**
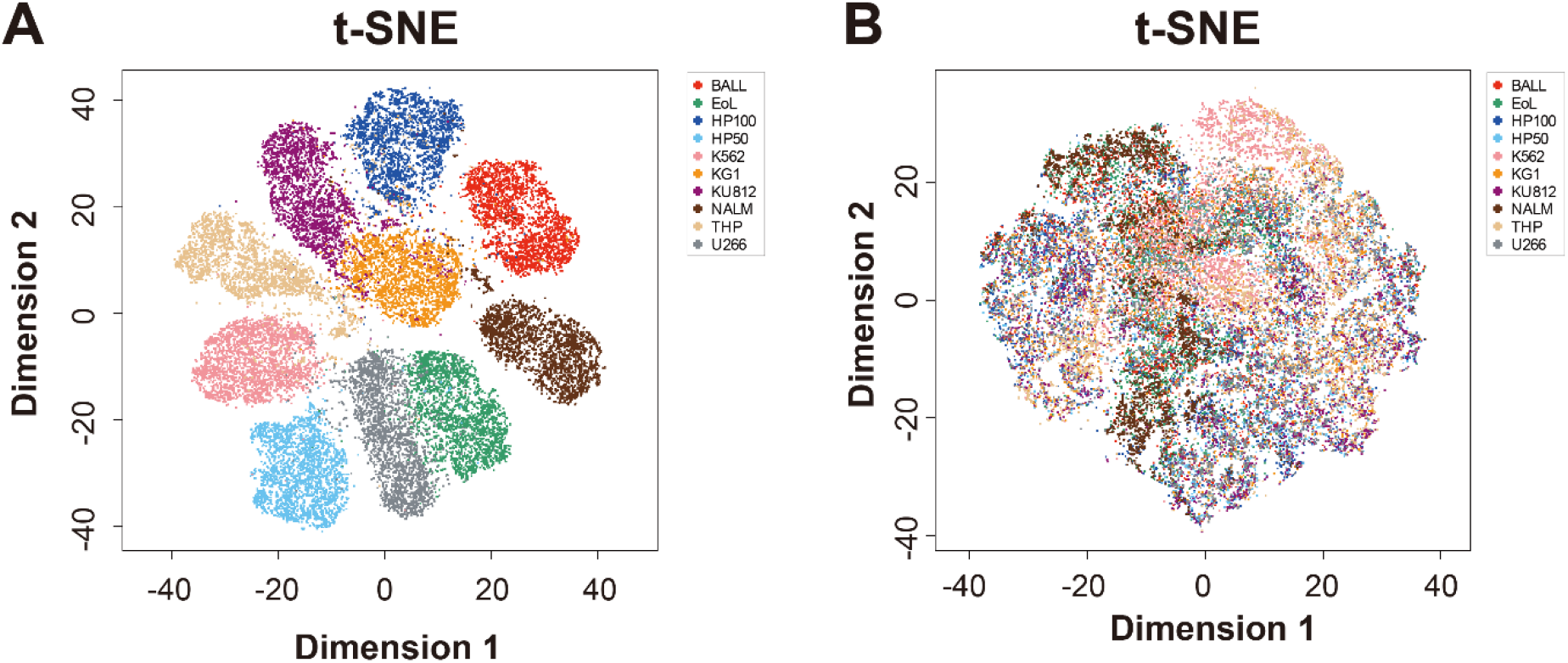
**Comparison of extracted features of each cell line by t-SNE** (A) The extracted features (128 features/image) and (B) the raw-digital data (2,304 values) were compressed into 2D by t-SNE and plotted in the scatter plot. The plotted data in (A) and (B) were obtained from the same training image (3,000 images/group).

### Concluding remarks

Collating the obtained findings together, the combination of IFC and CNN is a powerful tool for cell-classification and easily processes a large number of samples with high discrimination accuracy. In addition, this system has the merit that once the models of features of each cell population have been established, they can be readily used for subsequent analysis. Moreover, a more robust discrimination system could be established by adding images acquired in a later experiment. It is suggested that the issue of how to collect a huge number of images for model training is key for establishing a high-performance system for discriminating cell populations using CNN. Application of IFC for image acquisition of floating cells is expected to be employed in ML.

Currently, ML is also expected to be applicable to pathological diagnosis (Kourou et al., 2015; Doan M et al., 2018). It has been reported that the discrimination of skin cancers can be performed with dermatologist-level accuracy using CNN (Esteva et al., 2017). Generally, images of pathological regions are used for CNN. However, the system established in this study uses images of single cells to train a model for CNN. Therefore, it is theoretically possible to obtain images and features of over 1.0 × 10^6^ single hematopoietic cells from one hematopoietic tumor sample using a combination of IFC and CNN. This novel cell discrimination method could be widely applied to research in the field of cell biology as well as to diagnoses of clinical hematopoietic tumors.

## Materials and methods

### Reagents

All hematopoietic tumor cell lines were obtained from RIKEN BioResource Research Center and American Type Culture Collection. The details of each cell line are listed in Table S1. The cells were cultured in RPMI1640 (Nacalai Tesque, Kyoto, Japan) plus 10% fetal calf serum (Biofill, Elsternwick, Victoria, Australia) supplemented with 1× Non-Essential Amino Acid Solution (Thermo Fisher Scientific K.K., Tokyo, Japan), and 1 mM sodium pyruvate (Thermo Fisher Scientific K.K.) until they reached semi-confluence.

### Image recording

The bright-field images of single cells from each cell line were recorded using an Imaging Flow Cytometer ImageStreamX Mark II (Merck Ltd., Tokyo, Japan) with 40× objective lens without Extended Depth of Field option. Cultured cells were harvested and stained with 1:10,000 diluted DRAQ5 (BioLegend, San Diego, CA, USA) and 1:25 diluted FITC-conjugated Annexin V (Biolegend) in Annexin V Binding Buffer (Biolegend). Then, the cells were washed once with Annexin V Binding Buffer. The stained cells were re-suspended into 100 µL of Annexin V Binding Buffer. To normalize the bias regarding the focus and brightness among the experimental days, the cell images were separately recorded for three experimental days. Over 5.0 × 10^4^ bright-field images of each cell line were recorded at once.

### Processing of cell images of each cell line

The recorded cell images were exported in 8-bit grayscale TIFF format using IDEAS software (version 6.2). The images not including cells in the field and extremely deformed cells were manually excluded from the analyses. The randomly chosen individual 3.0 × 10^4^ and 1.0 × 10^3^ images/group obtained from each experimental day were combined. Finally, we obtained 9.0 × 10^5^ training data and 3.0 × 10^4^ test data set evenly including the 10 classes of cell lines for CNN as shown in Figure 3A. For discrimination of the three classes, training and test data were extracted from the data of 10 classes.

### CNN training

CNN was performed by Keras (version 2.1.5) with tensorflow-gpu (version 1.2.1) backend. The CNN was constructed with reference to the Keras official website (https://keras.io/). Input: 48 × 48 = 2304 data. Extracted features: 128 features/image.

The classification was performed by softmax using the extracted features.

### Machine spec

CNN was performed on a local machine and a virtual machine built on the Amazon Elastic Compute Cloud (Amazon EC2) as follows:

Local machine: OS Windows 7, GPU NVIDIA Quadro K620 with CUDA Toolkit (version 8.0) and cuDNN (version 5.1), Python 3.5 with Jupyter Notebook.

Amazon EC2: p2.xlarge instants, OS Windows Server 2016, GPU NVIDIA Tesla K80 with CUDA Toolkit (version 8.0) and cuDNN (version 5.1), Python 3.5 with Jupyter Notebook.

### t-Distributed Stochastic Neighbor Embedding (t-SNE)

t-SNE was performed using extracted features (128 features/image) and raw digital images (2,304 data) of each set of 3,000 images/group by R (version 3.4.1) with the Rtsne (version 0.13) package. These data were compressed into 2D and visualized in a scatter plot.

### Classification of each cell line by biologists

To test the classification accuracy for 10 classes of hematopoietic tumor cell lines, two biologists were trained using 140 images of each class of cell line as shown in Figure S1. For each test, 100 randomly chosen images generated from the data for model training of CNN were used. Each biologist classified the cells by referring to the training data. The test was repeated three times with the test data being changed each time.

### Data availability

Python code for CNN, csv format data for model training and variation, and trained model were deposited in the website (https://figshare.com/s/0ff584cb070cd03164aa).

## Author contributions

Y.M. designed and performed the experiments, analyzed and interpreted the data, provided financial and administrative support, and wrote the paper; R.N., F.T. performed the experiments, interpreted the data, and contributed to manuscript writing.

## Acknowledgments

We thank H. Gonda for the advice on using the image cytometer.

Funding: This work was supported by JSPS KAKENHI Grant Number JP17K09962.

